## Supplementary Materials for Local nutrient addition drives plant biodiversity losses but not biotic homogenization in global grasslands for "Local nutrient addition drives plant diversity losses but not biotic homogenization in global grasslands"

**The PDF file includes:**

Figs. S1 to S10

Tables S1 to S8

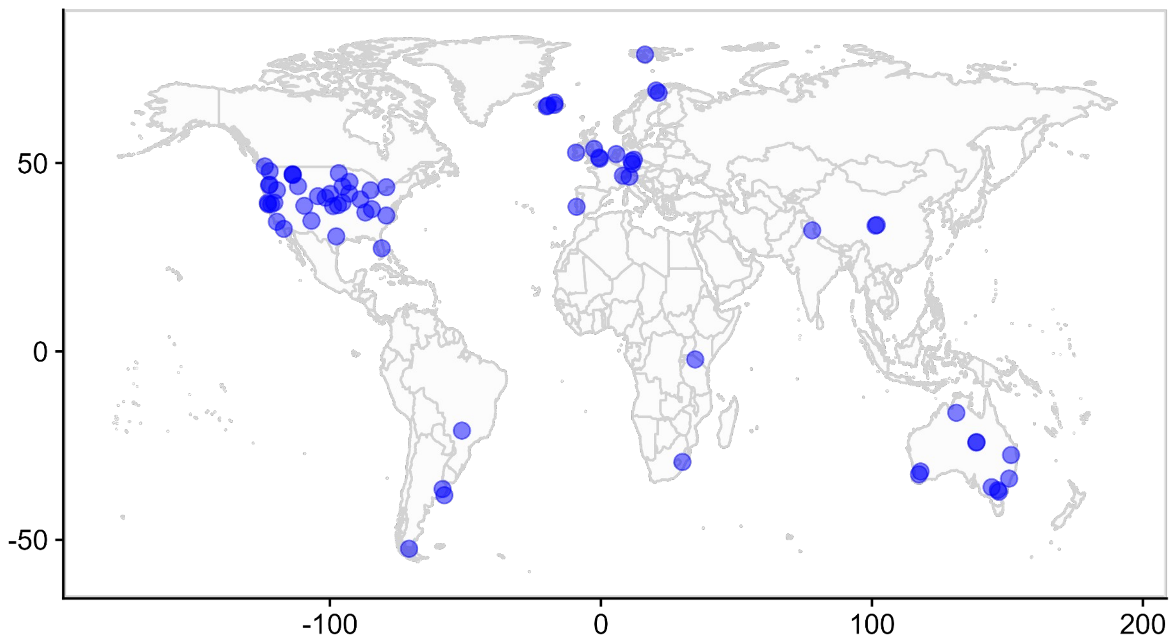


**Fig. S1.**

**Geolocation of the 72 sites used in this analysis.** See table S1 for more detail about site information such as habitat type, experimental years. Source data are provided as a Source Data file.

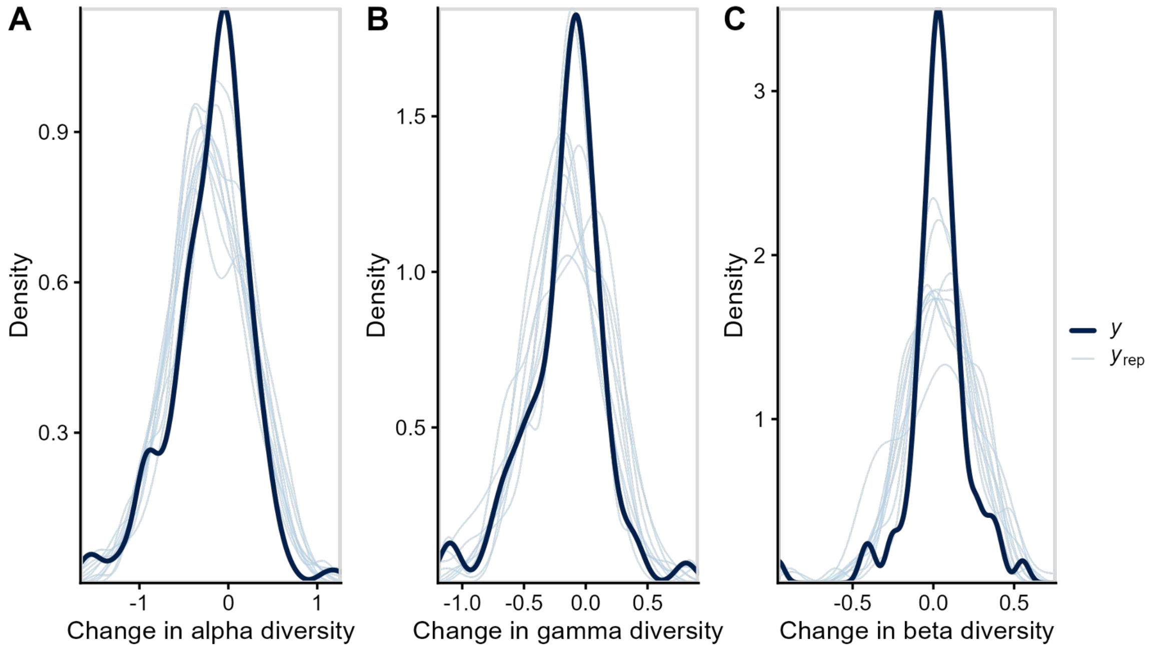


**Fig. S2.**

**Predicted values (the thin lines) and observed values (the thick lines) for change in alpha, gamma, and beta diversity (Δα, Δγ, and Δβ) under nutrient addition.** Model predicted values reasonably well. See Table S3 for more detail about model fitting.

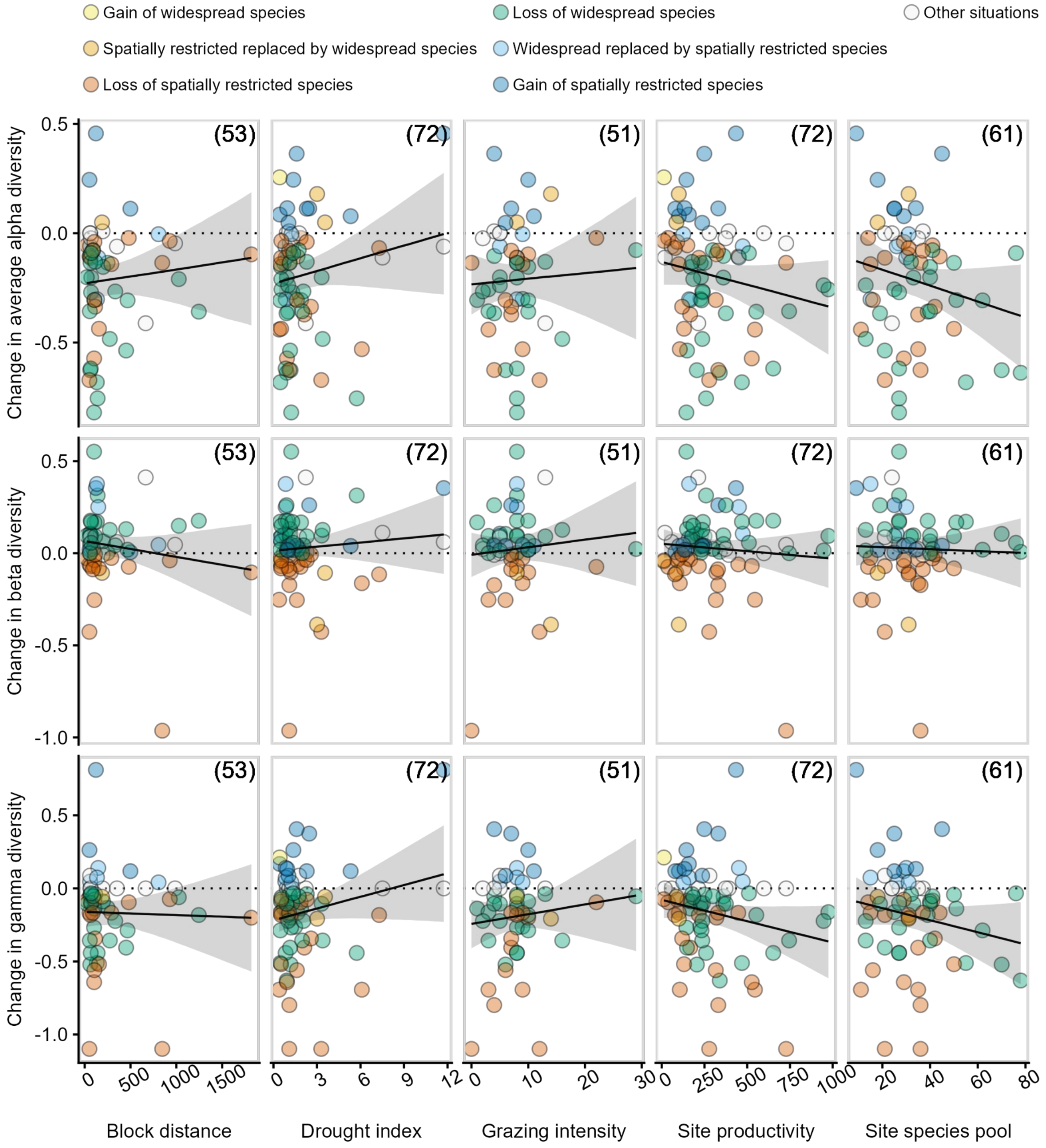


**Fig. S3.**

**Bivariate relationship between change in average alpha, gamma, and beta diversity ( $\overline{\Delta\alpha}$ , Δγ, and Δβ) 4 years after treatment began and site covariates.** Unit for block distance is meter, unit for site productivity is g $m^{-2}$. Solid black lines represent linear regression lines, grey areas are 95% credible bands. None of the relationships were statistically significant. Numbers in the parentheses are the number of sites. Other scenarios: $\overline{\Delta\alpha}$ = 0, Δγ = 0, or Δβ = 0, when a site has these situations, it was not counted into any of the six scenarios in the framework. Site sier.us was excluded for the relationship between block distance and change in diversity because blocks were arranged very far away from each other (12538.09 m), $\overline{\Delta\alpha}$ , Δγ, and Δβ was 0.364, 0.405, and 0.041, respectively. Source data are provided as a Source Data file.

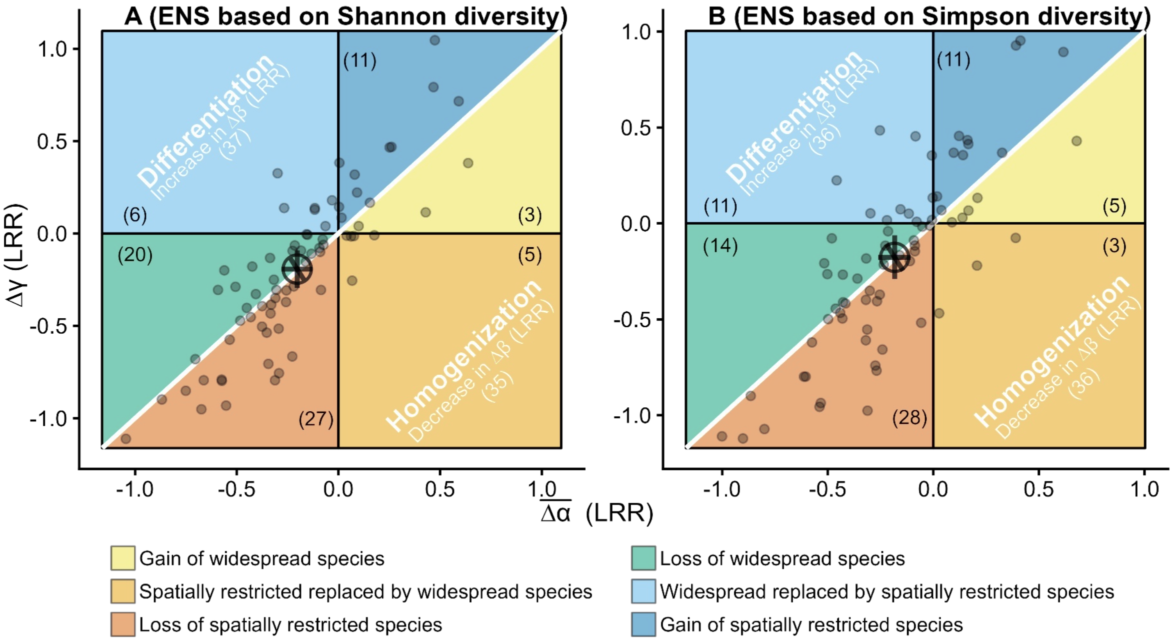


**Fig. S4.**

**Changes in average α, γ, and β diversity ( $\overline{\Delta\alpha}$ , Δγ, and Δβ) with nutrient addition using effective number of species based on (A) Shannon diversity (B) Simpson diversity**. ENS: effective number of species; LRR: log response ratio. The white 1:1 diagonal line indicates no effects of nutrient addition on β diversity. Numbers in the parentheses are the number of sites. When a site has $\overline{\Delta\alpha}$ = 0, Δγ = 0, or Δβ = 0, it was not counted into any of the six scenarios in the key. The small points represent site-level $\overline{\Delta\alpha}$ , Δγ, and Δβ at 72 sites. The large open point and error bars are the estimated mean and 95% credible intervals for $\overline{\Delta\alpha}$ , Δγ, and Δβ across all sites. See Table S3 for model fit and estimated overall means and 95% credible intervals for $\overline{\Delta\alpha}$ , Δγ, and Δβ. Source data are provided as a Source Data file.

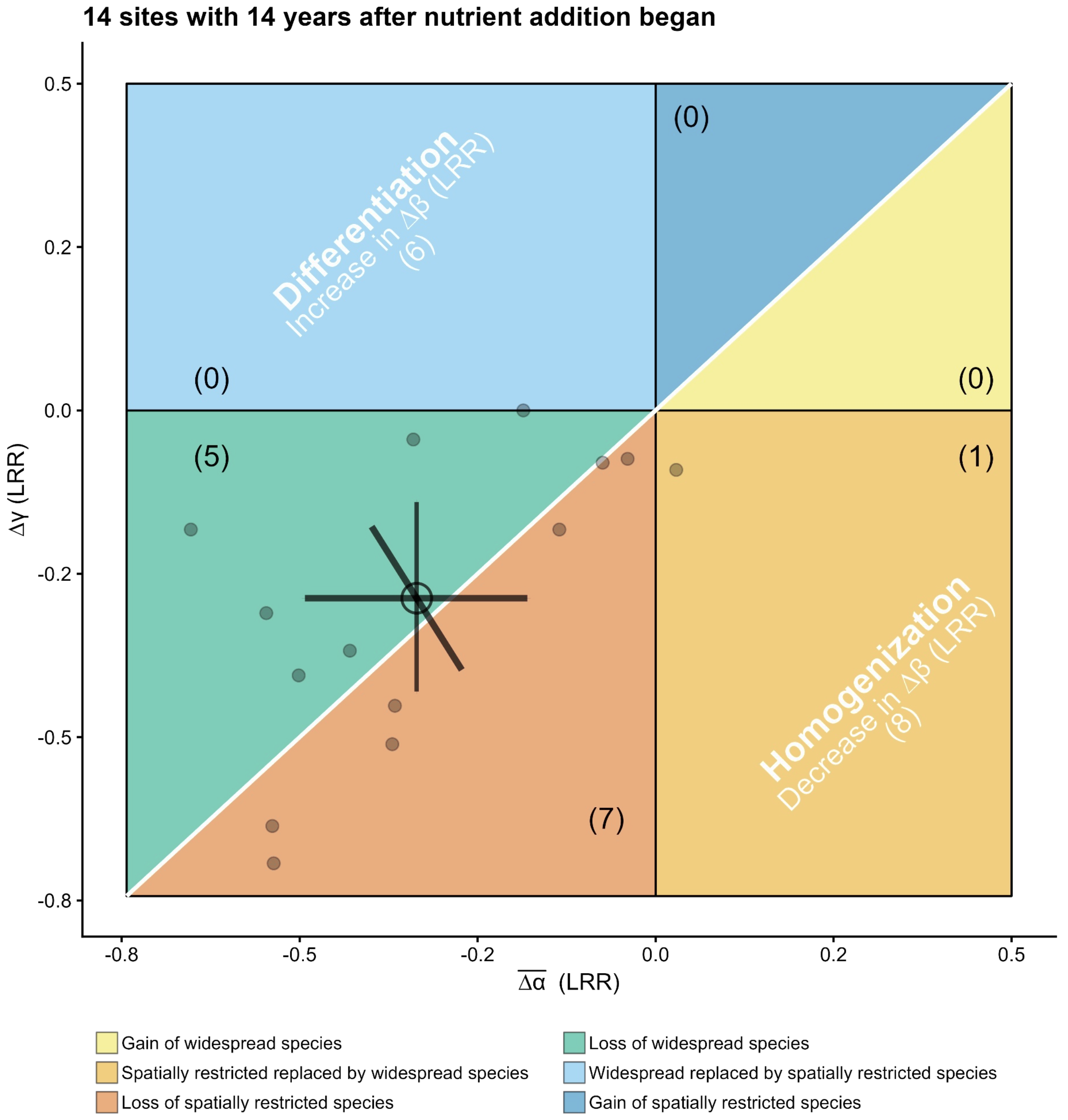


**Fig. S5.**

**Changes in average α, γ, and β diversity ( $\overline{\Delta\alpha}$ , Δγ, and Δβ) with nutrient addition 14 years after treatment began.** LRR: log response ratio. The white 1:1 diagonal line indicates no effects of nutrient addition on β diversity. Numbers in the parentheses are the number of sites. When a site has $\overline{\Delta\alpha}$ = 0, Δγ = 0, or Δβ = 0, it was not counted into any of the six scenarios. The small points represent site-level $\overline{\Delta\alpha}$ , Δγ, and Δβ. The large open point and error bars are the estimated mean and 95% credible intervals for $\overline{\Delta\alpha}$ , Δγ, and Δβ across all sites. Source Data are provided as a Source Data file.

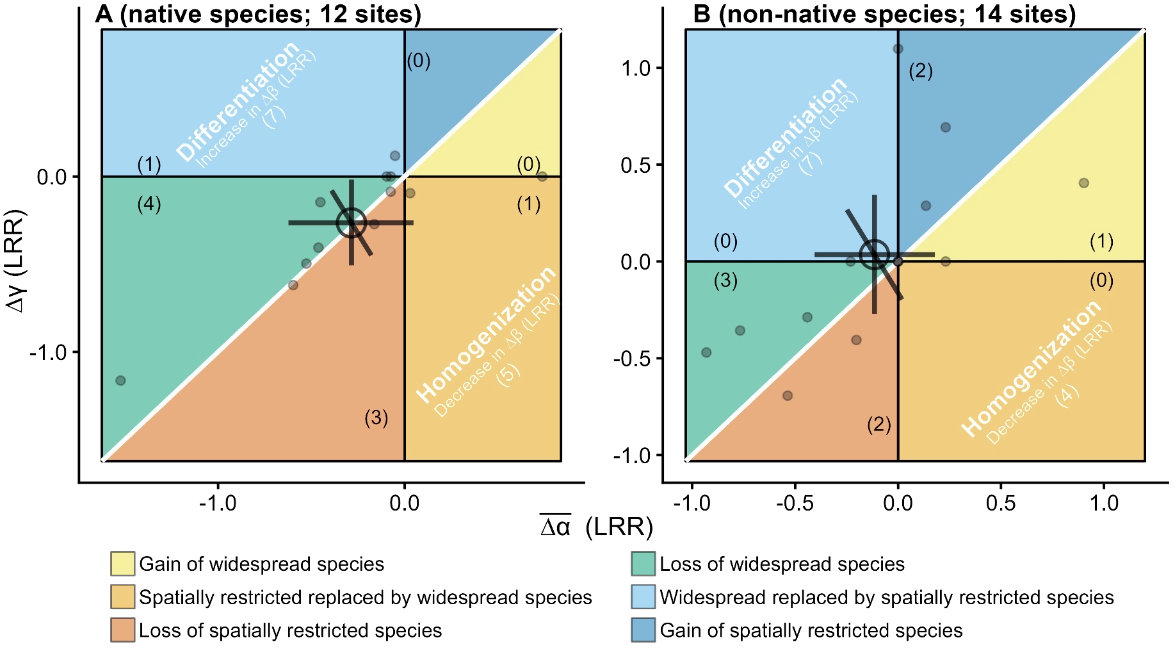


**Fig. S6.**

**Changes in average α, γ, and β diversity ( $\overline{\Delta\alpha}$ , Δγ, and Δβ) with nutrient addition for native and non-native species groups 14 years after treatment began. (A) native and (B) non-native species.** LRR: log response ratio. The white 1:1 diagonal line indicates no effects of nutrient addition on β diversity. Numbers in the parentheses are the number of sites. When a site has $\overline{\Delta\alpha}$ = 0, Δγ = 0, or Δβ = 0, it was not counted into any of the six scenarios. The small points represent site-level $\overline{\Delta\alpha}$ , Δγ, and Δβ. The large open point and error bars are the estimated mean and 95% credible intervals for $\overline{\Delta\alpha}$ , Δγ, and Δβ across all sites. Source data are provided as a Source Data file.

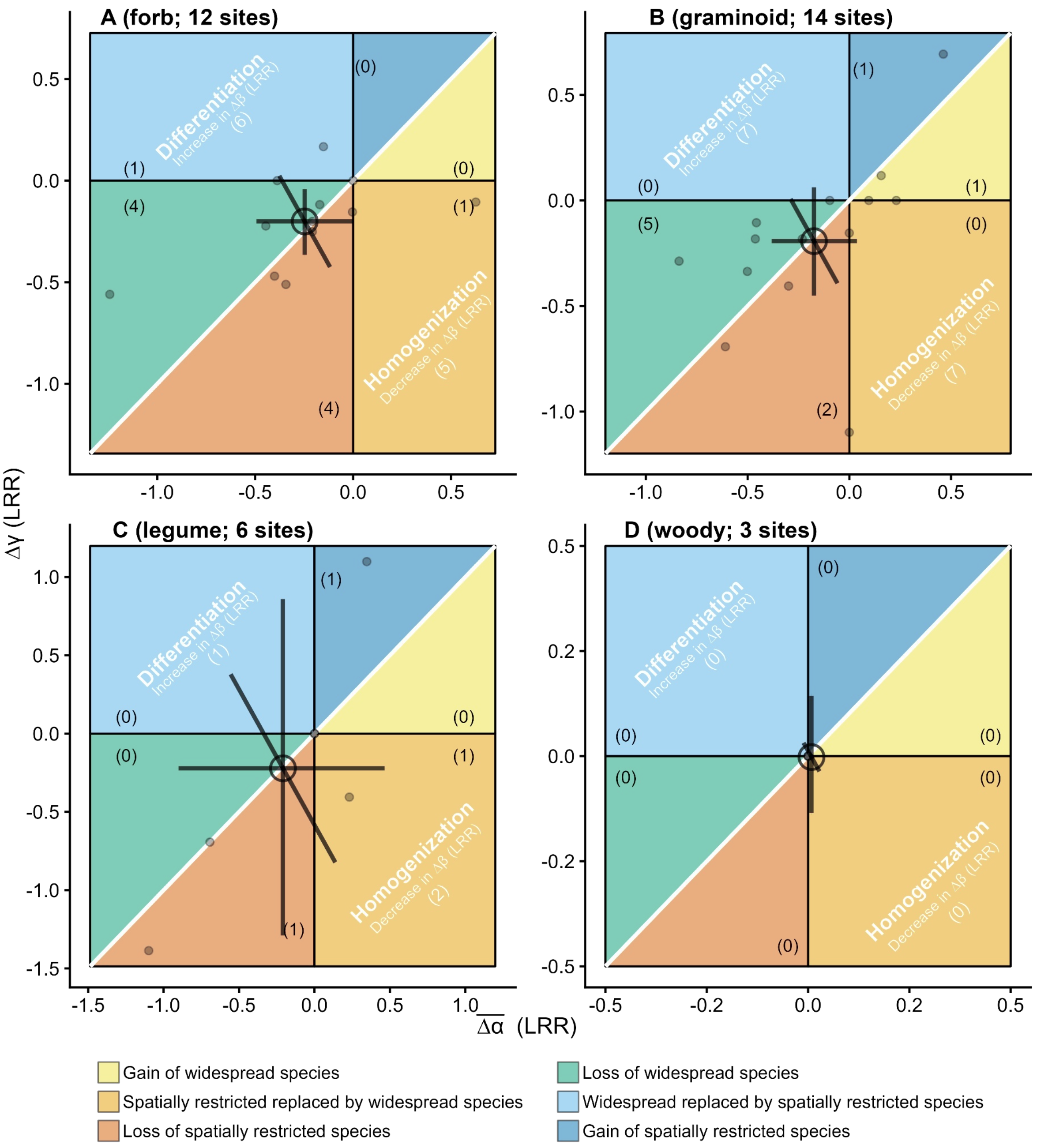


**Fig. S7.**

**Changes in average α, γ, and β diversity ( $\overline{\Delta\alpha}$ , Δγ, and Δβ) with nutrient addition for different functional species groups 14 years after treatment began. (A) forb, (B) graminoid, (C) legume, and (D) woody species.** LRR: log response ratio. The white 1:1 diagonal line indicates no effects of nutrient addition on β diversity. Numbers in the parentheses are the number of sites. When a site has $\overline{\Delta\alpha}$ = 0, Δγ = 0, or Δβ = 0, it was not counted into any of the six scenarios. The small points represent site-level $\overline{\Delta\alpha}$ , Δγ, and Δβ. The large open point and error bars are the estimated mean and 95% credible intervals for $\overline{\Delta\alpha}$ , Δγ, and Δβ across all sites. Source data are provided as a Source Data file.

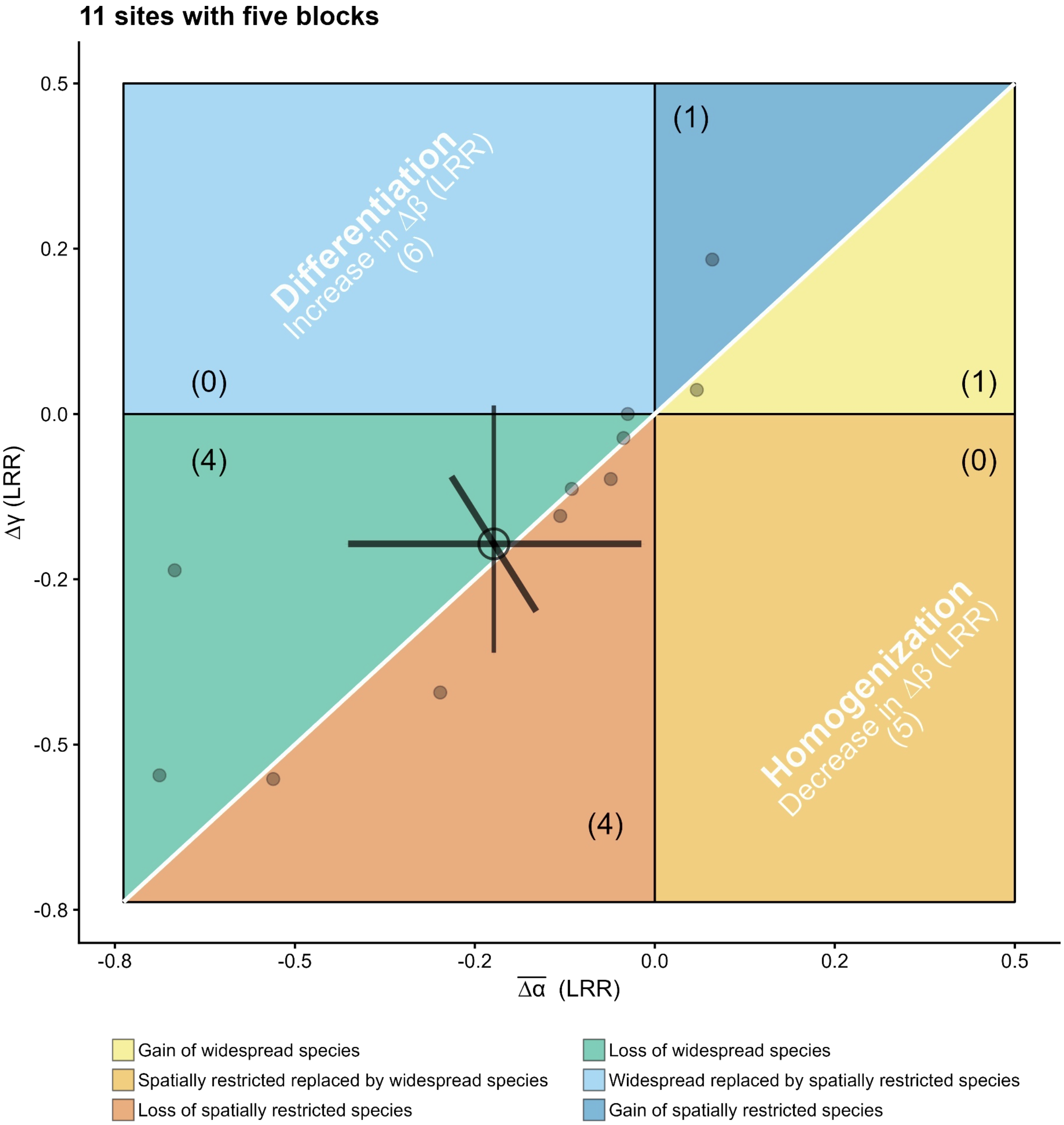


**Fig. S8.**

**Changes in average α, γ, and β diversity ( $\overline{\Delta\alpha}$ , Δγ, and Δβ) with nutrient addition for sites with 5 blocks.** LRR: log response ratio. The white 1:1 diagonal line indicates no effects of nutrient addition on β diversity. Numbers in the parentheses are the number of sites. When a site has $\overline{\Delta\alpha}$ = 0, Δγ = 0, or Δβ = 0, it was not counted into any of the six scenarios. The small points represent site-level $\overline{\Delta\alpha}$ , Δγ, and Δβ. The large open point and error bars are the estimated mean and 95% credible intervals for $\overline{\Delta\alpha}$ , Δγ, and Δβ across all sites. Source data are provided as a Source Data file.

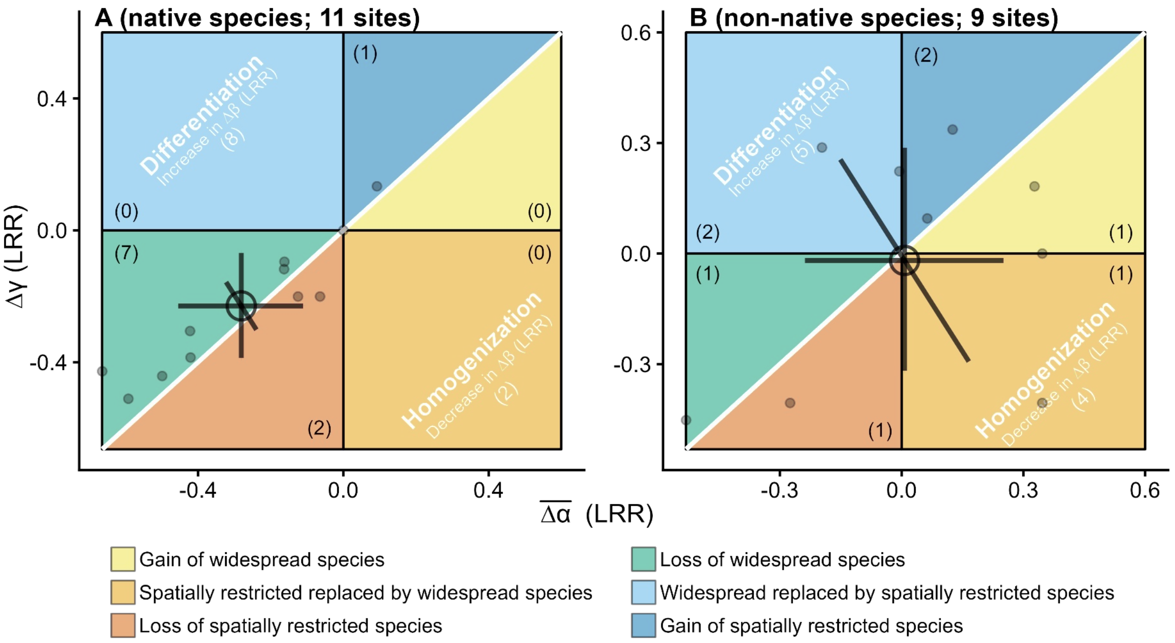


**Fig. S9.**

Changes in average α, γ, and β diversity ( $\overline{\Delta\alpha}$ , Δγ, and Δβ) with nutrient addition for native and non-native species groups for 11 sites with 5 blocks. (A) native and (B) non-native species. LRR: log response ratio. The white 1:1 diagonal line indicates no effects of nutrient addition on β diversity. Numbers in the parentheses are the number of sites. When a site has $\overline{\Delta\alpha}$ = 0, Δγ = 0, or Δβ = 0, it was not counted into any of the six scenarios. The small points represent site-level $\overline{\Delta\alpha}$ , Δγ, and Δβ. The large open point and error bars are the estimated mean and 95% credible intervals for $\overline{\Delta\alpha}$ , Δγ, and Δβ across all sites. Source data are provided as a Source Data file.

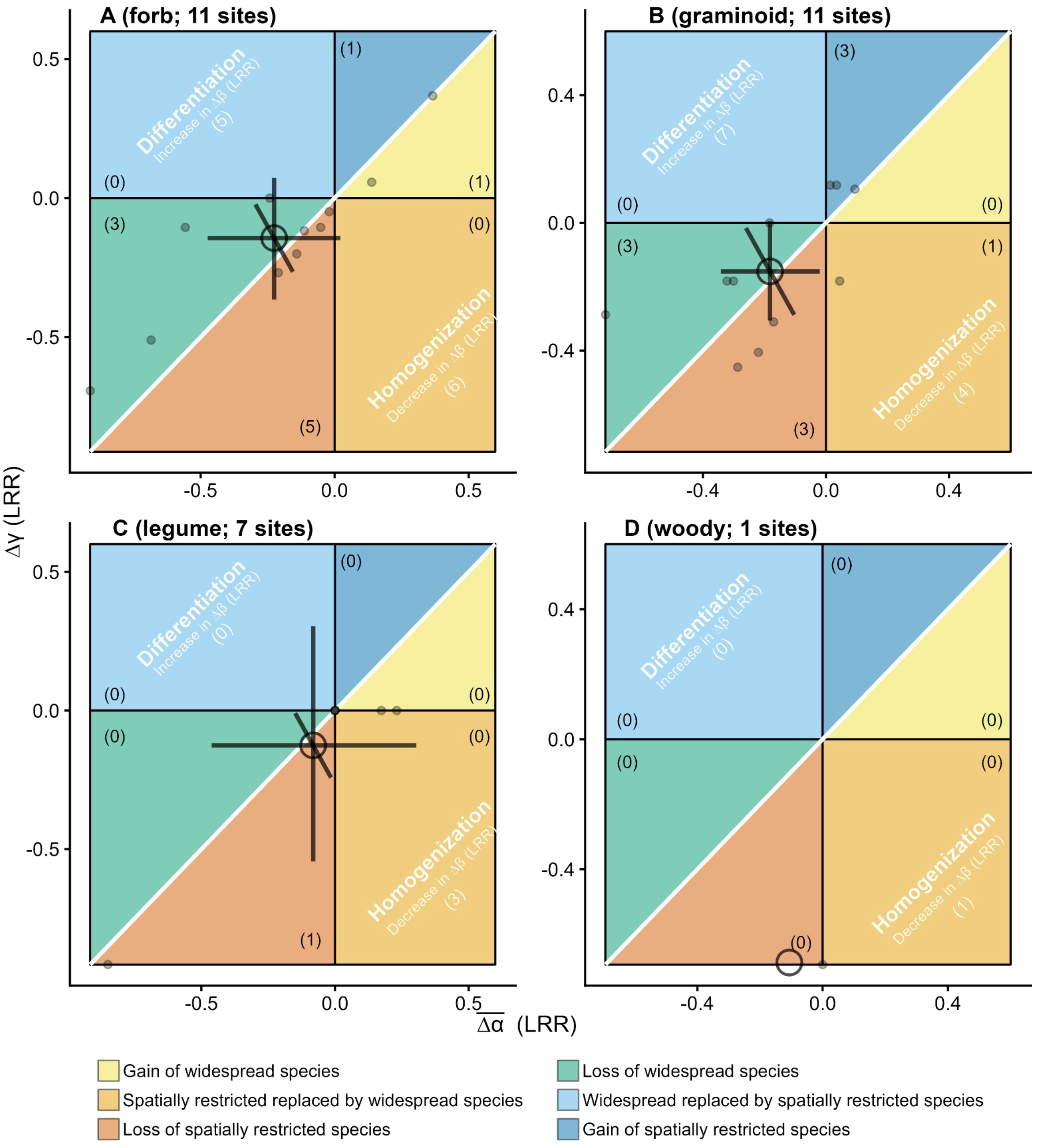


**Fig. S10.**

**Changes in average α, γ, and β diversity ( $\overline{\Delta\alpha}$ , Δγ, and Δβ) with nutrient addition for different functional species groups for 11 sites with 5 blocks. (A) forb, (B) graminoid, (C) legume, and (D) woody species.** LRR: log response ratio. The white 1:1 diagonal line indicates no effects of nutrient addition on β diversity. Numbers in the parentheses are the number of sites. When a site has $\overline{\Delta\alpha}$ = 0, Δγ = 0, or Δβ = 0, it was not counted into any of the six scenarios. The small points represent site-level $\overline{\Delta\alpha}$ , Δγ, and Δβ. The large open point and error bars are the estimated mean and 95% credible intervals for $\overline{\Delta\alpha}$ , Δγ, and Δβ across all sites. Source data are provided as a Source Data file.

**Table S1.**
Site geolocation, grassland types, and experimental years used.

| site_code | Continent | Habitat | Latitude | Longitude | First nutrient year | Year used |
|---|---|---|---|---|---|---|
| ahth.is | Europe | heathland | 65.133350 | -19.672060 | 2016 | 4 |
| amlr.is | Europe | desert grassland | 65.133350 | -19.672060 | 2016 | 4 |
| arch.us | North America | mixedgrass prairie | 27.170340 | -81.218280 | 2016 | 4 |
| azi.cn | Asia | alpine grassland | 33.670000 | 101.870000 | 2008 | 4 |
| azitwo.cn | Asia | alpine grassland | 33.580000 | 101.530000 | 2019 | 4 |
| badlau.de | Europe | old field | 51.392138 | 11.877969 | 2016 | 4 |
| barta.us | North America | mixedgrass prairie | 42.244586 | -99.651800 | 2008 | 4 |
| bayr.de | Europe | mesic grassland | 49.916667 | 11.583333 | 2017 | 4 |
| bnbt.us | North America | tallgrass prairie | 39.597007 | -95.086276 | 2018 | 4 |
| bnch.us | North America | montane grassland | 44.276685 | -121.968017 | 2008 | 4, 14 |
| bogong.au | Australia | alpine grassland | -36.874000 | 147.254000 | 2010 | 4, 14 |
| burrawan.au | Australia | semiarid grassland | -27.736138 | 151.139131 | 2009 | 4 |
| burren.ie | Europe | calcareous grassland | 53.072023 | -8.992624 | 2016 | 4 |
| cbgb.us | North America | tallgrass prairie | 41.785067 | -93.385383 | 2010 | 4 |
| cdcr.us | North America | tallgrass prairie | 45.425000 | -93.211500 | 2008 | 4, 14 |
| cdpt.us | North America | shortgrass prairie | 41.206000 | -101.643000 | 2008 | 4, 14 |
| chilcas.ar | South America | mesic grassland | -36.275556 | -58.265556 | 2014 | 4 |
| comp.pt | Europe | annual grassland | 38.829280 | -8.791406 | 2013 | 4 |
| cowi.ca | North America | old field | 48.808800 | -123.630100 | 2008 | 4, 14 |
| elliot.us | North America | annual grassland | 32.875000 | -117.052243 | 2009 | 4, 14 |
| ethamc.au | Australia | desert grassland | -23.756333 | 138.474283 | 2014 | 4 |

| site_code | Continent | Habitat | Latitude | Longitude | First nutrient year | Year used |
|---|---|---|---|---|---|---|
| ethass.au | Australia | desert grassland | -23.638610 | 138.398970 | 2014 | 4 |
| frue.ch | Europe | pasture | 47.114227 | 8.540831 | 2009 | 4 |
| hall.us | North America | tallgrass prairie | 36.871944 | -86.701670 | 2008 | 4 |
| hart.us | North America | shrub steppe | 42.723745 | -119.497670 | 2008 | 4 |
| hero.uk | Europe | mesic grassland | 51.411000 | -0.639000 | 2008 | 4, 14 |
| hopl.us | North America | annual grassland | 39.012753 | -123.060313 | 2008 | 4 |
| jena.de | Europe | grassland | 50.938000 | 11.528347 | 2014 | 4 |
| kbs.us | North America | old field | 42.408986 | -85.390900 | 2014 | 4 |
| kibber.in | Asia | alpine grassland | 32.319727 | 78.009540 | 2012 | 4 |
| kidman.au | Australia | savanna | -16.108460 | 130.953100 | 2015 | 4 |
| kilp.fi | Europe | tundra grassland | 69.056700 | 20.874700 | 2014 | 4 |
| kiny.au | Australia | semiarid grassland | -36.200000 | 143.750000 | 2008 | 4, 14 |
| koffler.ca | North America | pasture | 44.024047 | -79.535881 | 2011 | 4 |
| konz.us | North America | tallgrass prairie | 39.070856 | -96.582821 | 2008 | 4, 14 |
| lagoas.br | South America | cerrado | -20.982778 | -51.798889 | 2017 | 4 |
| lake.us | North America | tallgrass prairie | 43.384683 | -95.181052 | 2016 | 4 |
| lancaster.uk | Europe | mesic grassland | 53.985625 | -2.628418 | 2009 | 4 |
| look.us | North America | montane grassland | 44.205177 | -122.128447 | 2008 | 4, 14 |
| marc.ar | South America | grassland | -37.715160 | -57.424510 | 2012 | 4 |
| mcla.us | North America | annual grassland | 38.864272 | -122.406406 | 2008 | 4 |
| moab.us | North America | NA | 38.782000 | -109.652000 | 2019 | 4 |
| msla.us | North America | grassland | 46.664613 | -114.000523 | 2018 | 4 |
| msla_2.us | North | grassland | 46.664887 | -114.001321 | 2018 | 4 |

| site_code | Continent | Habitat | Latitude | Longitude | First nutrient year | Year used |
|---|---|---|---|---|---|---|
| | America | | | | | |
| msla_3.us | North America | grassland | 46.664744 | -114.001678 | 2018 | 4 |
| msum.us | North America | tallgrass prairie | 46.869624 | -96.453523 | 2017 | 4 |
| mtca.au | Australia | savanna | -31.782138 | 117.610853 | 2009 | 4 |
| nilla.au | Australia | old field | -36.896000 | 146.007000 | 2017 | 4 |
| ping.au | Australia | old field | -32.496169 | 116.972947 | 2014 | 4 |
| potrok.ar | South America | semiarid grassland | -51.916028 | -70.407444 | 2016 | 4 |
| rook.uk | Europe | mesic grassland | 51.406000 | -0.644000 | 2008 | 4 |
| saana.fi | Europe | montane grassland | 69.036800 | 20.843300 | 2015 | 4 |
| sage.us | North America | montane grassland | 39.430000 | -120.240000 | 2008 | 4 |
| saline.us | North America | mixedgrass prairie | 39.050000 | -99.100000 | 2008 | 4 |
| sedg.us | North America | annual grassland | 34.700000 | -120.016667 | 2008 | 4 |
| sereng.tz | Africa | savanna | -2.429387 | 34.857435 | 2009 | 4 |
| sevi.us | North America | desert grassland | 34.359200 | -106.690500 | 2008 | 4, 14 |
| sgs.us | North America | shortgrass prairie | 40.816670 | -104.766670 | 2008 | 4, 14 |
| shps.us | North America | shrub steppe | 44.257212 | -112.205337 | 2008 | 4, 14 |
| sier.us | North America | annual grassland | 39.235510 | -121.283696 | 2008 | 4 |
| smith.us | North America | mesic grassland | 48.206581 | -122.624754 | 2008 | 4 |
| spin.us | North America | pasture | 38.132222 | -84.502778 | 2008 | 4, 14 |
| sval.no | Europe | tundra grassland | 78.690180 | 16.447120 | 2019 | 4 |
| temple.us | North America | tallgrass prairie | 31.044000 | -97.349000 | 2008 | 4, 14 |
| thth.is | Europe | heathland | 65.901290 | -17.089270 | 2016 | 4 |
| tmlr.is | Europe | desert grassland | 65.901290 | -17.089270 | 2016 | 4 |
| trel.us | North America | tallgrass prairie | 40.075000 | -88.829000 | 2009 | 4 |

| site_code | Continent | Habitat | Latitude | Longitude | First nutrient year | Year used |
|---|---|---|---|---|---|---|
| ukul.za | Africa | mesic grassland | -29.670238 | 30.401051 | 2010 | 4 |
| unc.us | North America | old field | 36.008280 | -79.020423 | 2008 | 4 |
| valm.ch | Europe | alpine grassland | 46.631345 | 10.372252 | 2009 | 4 |
| veluwe.nl | Europe | old field | 52.026547 | 5.807947 | 2018 | 4 |
| yarra.au | Australia | mesic grassland | -33.613694 | 150.738031 | 2015 | 4 |

**Table S2.**
**Summary of spatial autocorrelation for the effects of nutrient addition on alpha, gamma, and beta diversity (Δα, Δγ, and Δβ) four year after nutrient addition began using Moran's I test.** Positive Moran's I suggests similar values cluster together while negative Moran's I suggests dissimilar values are adjacent. The z-score quantifies how extreme the observed Moran's I value is compared with that is expected under the null hypothesis of spatial randomness.

| Diversity facets | Moran's I | Standard deviate (z score) | p |
|---|---|---|---|
| Δα | 0.1258 | 1.2920 | 0.0982 |
| Δγ | 0.1274 | 1.3251 | 0.0926 |
| Δβ | -0.0312 | -0.1687 | 0.5670 |

**Table S3.**

**Summary of model fitting for the effects of nutrient addition on alpha, gamma, and beta diversity (Δα, Δγ, and Δβ) four years after treatment began.** We calculated α and γ diversity using species richness, effective number of species (ENS) based on Shannon diversity, and Simpson diversity. We fitted the random intercept models using the brm function. For Δα, Δγ, and Δβ, we included random intercepts among sites, model was coded as: diversity (log scale) ~ 1 + (1 |sites). Bulk-ESS is the effective sample size, is useful as a diagnostic for the sampling efficiency. Tail_ESS is the minimum of the effective sample sizes for 5% and 95% quantiles. Low Rhats indicates model convergence were achieved. All models used 1000 iterations for warmup and 3000 iterations

| Diversity metrics | Diversity facets | Estimate | l-95% CI | u-95% CI | Rhat | Bulk_ESS | Tail_ESS | Number of sites |
|---|---|---|---|---|---|---|---|---|
| Species richness | Δα | -0.1916 | -0.2549 | -0.1297 | 1.0013 | 16067.290 | 9466.034 | 72 |
| Species richness | Δβ | 0.0280 | -0.0208 | 0.0763 | 1.0027 | 2133.078 | 1718.600 | 72 |
| Species richness | Δγ | -0.1623 | -0.2365 | -0.0848 | 1.0040 | 1583.065 | 1456.179 | 72 |
| Simpson diversity | Δα | -0.1833 | -0.2611 | -0.1062 | 1.0006 | 11229.847 | 9428.506 | 72 |
| Simpson diversity | Δβ | 0.0083 | -0.0599 | 0.0767 | 1.0013 | 3977.223 | 5845.292 | 72 |
| Simpson diversity | Δγ | -0.1768 | -0.2896 | -0.0616 | 1.0019 | 2636.618 | 5521.383 | 72 |
| Shannon diversity | Δα | -0.2027 | -0.2802 | -0.1269 | 1.0001 | 7935.751 | 8781.933 | 72 |
| Shannon diversity | Δβ | 0.0102 | -0.0423 | 0.0632 | 1.0028 | 2269.188 | 3430.052 | 72 |
| Shannon diversity | Δγ | -0.1933 | -0.2953 | -0.0919 | 1.0022 | 2596.788 | 3157.553 | 72 |

**Table S4.**

**Estimated mean and 95% credible intervals for change in alpha, gamma, and beta diversity ( $\overline{\Delta\alpha}$ , Δγ, and Δβ) at individual sites four years under nutrient addition began.** Diversity was measured as species richness. Scenarios differ from that described in the main text because here we considered the 95% credible intervals estimated from models. Other situation refers to $\overline{\Delta\alpha}$ overlapped with 0, or Δγ overlapped with 0, or $\overline{\Delta\alpha}$ overlapped with 0, Δγ overlapped with 0, and Δβ overlapped with 0, when a site has these situations, it was not counted into any of the six scenarios in the framework.

| site_code | Δα [l-95% CI, u-95% CI] | Δγ [l-95% CI, u-95% CI] | Δβ [l-95% CI, u-95% CI] | Change in widespread or spatially restricted species |
|---|---|---|---|---|
| ahth.is | -0.1513 [-0.3345, 0.0898] | -0.0018 [-0.2977, 0.2935] | 0.0524 [-0.1133, 0.2255] | Other situations |
| amlr.is | -0.1293 [-0.3044, 0.1305] | 0.0195 [-0.2862, 0.3364] | -0.004 [-0.1752, 0.1597] | Other situations |
| arch.us | -0.1937 [-0.394, 0.0081] | -0.1352 [-0.3858, 0.1129] | 0.0588 [-0.1064, 0.2306] | Other situations |
| azi.cn | -0.1797 [-0.3702, 0.0275] | -0.1103 [-0.3606, 0.145] | 0.038 [-0.1251, 0.2042] | Other situations |
| azitwo.cn | -0.2559 [-0.5218, -0.0785] | -0.3898 [-0.7254, -0.0557] | 0.0193 [-0.1472, 0.1824] | Loss of spatially restricted species |
| badlau.de | -0.1815 [-0.3796, 0.0313] | -0.1661 [-0.413, 0.0768] | -0.0104 [-0.185, 0.154] | Other situations |
| barta.us | -0.2067 [-0.4218, -0.0162] | -0.2124 [-0.4626, 0.0391] | 0.0328 [-0.1301, 0.1974] | Other situations |
| bayr.de | -0.1721 [-0.3595, 0.0459] | -0.1245 [-0.3836, 0.1291] | 0.0041 [-0.1662, 0.167] | Other situations |
| bnbt.us | -0.171 [-0.3579, 0.0509] | -0.0852 [-0.3386, 0.183] | 0.0357 [-0.1256, 0.2019] | Other situations |
| bnch.us | -0.1835 [-0.3783, 0.0256] | -0.1495 [-0.4095, 0.1021] | 0.0171 [-0.1529, 0.1817] | Other situations |
| bogong.au | -0.1724 [-0.3579, 0.0397] | -0.0607 [-0.3263, 0.2094] | 0.063 [-0.104, 0.2376] | Other situations |
| burrawan.au | -0.2235 [-0.4485, -0.0414] | -0.0855 [-0.3425, 0.1832] | 0.1973 [-0.0297, 0.4582] | Other situations |
| burren.ie | -0.2622 [-0.5309, -0.0873] | -0.3326 [-0.6436, -0.0315] | 0.0903 [-0.0789, 0.2777] | Loss of spatially restricted species |
| cbgb.us | -0.178 [-0.3726, 0.0382] | -0.1813 [-0.4297, 0.0712] | -0.0292 [-0.2092, 0.1388] | Other situations |
| cdcr.us | -0.2813 [-0.5773, -0.101] | -0.2129 [-0.467, 0.0364] | 0.2582 [-0.0123, 0.5784] | Other situations |
| cdpt.us | -0.2165 [-0.4429, -0.0294] | -0.2818 [-0.5598, -0.0101] | 0.0014 [-0.1685, 0.1683] | Loss of spatially restricted species |
| chilcas.ar | -0.183 [-0.3801, | -0.6158 [-1.1399, | -0.408 [-0.9449, | Other situations |

| site_code | Δα [l-95% CI, u-95% CI] | Δγ [l-95% CI, u-95% CI] | Δβ [l-95% CI, u-95% CI] | Change in widespread or spatially restricted species |
|---|---|---|---|---|
| | 0.0232] | -0.1047] | 0.0524] | |
| comp.pt | -0.2074 [-0.4253, -0.0128] | -0.1502 [-0.396, 0.0948] | 0.0906 [-0.0763, 0.2792] | Other situations |
| cowi.ca | -0.228 [-0.4582, -0.05] | -0.4199 [-0.7836, -0.0737] | -0.0956 [-0.3162, 0.0988] | Loss of spatially restricted species |
| elliot.us | -0.2713 [-0.5577, -0.0976] | -0.299 [-0.5896, -0.02] | 0.1546 [-0.0396, 0.3784] | Loss of spatially restricted species |
| ethamc.au | -0.1729 [-0.357, 0.0382] | -0.0851 [-0.3493, 0.1888] | 0.0415 [-0.1184, 0.2063] | Other situations |
| ethass.au | -0.099 [-0.2788, 0.2066] | 0.3106 [-0.2176, 0.8525] | 0.1722 [-0.032, 0.4095] | Other situations |
| frue.ch | -0.2021 [-0.4151, -0.008] | -0.1624 [-0.4027, 0.0876] | 0.057 [-0.106, 0.2292] | Other situations |
| hall.us | -0.1647 [-0.3506, 0.0631] | -0.0837 [-0.3424, 0.1758] | 0.0172 [-0.1511, 0.1812] | Other situations |
| hart.us | -0.1532 [-0.3296, 0.0811] | -0.0257 [-0.3082, 0.255] | 0.0329 [-0.1316, 0.2015] | Other situations |
| hero.uk | -0.1799 [-0.3744, 0.0321] | -0.0154 [-0.3022, 0.2698] | 0.1258 [-0.0578, 0.3283] | Other situations |
| hopl.us | -0.254 [-0.515, -0.0751] | -0.3385 [-0.6433, -0.0408] | 0.0625 [-0.1024, 0.2365] | Loss of spatially restricted species |
| jena.de | -0.165 [-0.3477, 0.0586] | -0.085 [-0.3421, 0.1809] | 0.0112 [-0.1609, 0.1715] | Other situations |
| kbs.us | -0.2452 [-0.4933, -0.0684] | -0.3971 [-0.7384, -0.0627] | -0.0141 [-0.1881, 0.1459] | Loss of spatially restricted species |
| kibber.in | -0.1677 [-0.3563, 0.0563] | -0.1305 [-0.3816, 0.1251] | -0.0166 [-0.1962, 0.1491] | Other situations |
| kidman.au | -0.148 [-0.3254, 0.0941] | 0.0958 [-0.2537, 0.4568] | 0.1309 [-0.0511, 0.3364] | Other situations |
| kilp.fi | -0.1849 [-0.3804, 0.0205] | -0.1291 [-0.3762, 0.1214] | 0.0446 [-0.1193, 0.214] | Other situations |
| kiny.au | -0.1923 [-0.3947, 0.0125] | -0.1329 [-0.3791, 0.1185] | 0.0582 [-0.1084, 0.2351] | Other situations |
| koffler.ca | -0.2522 [-0.5119, -0.0763] | -0.2996 [-0.5921, -0.0197] | 0.0935 [-0.0748, 0.2781] | Loss of spatially restricted species |
| konz.us | -0.1688 [-0.3615, 0.0577] | -0.0837 [-0.3409, 0.1872] | 0.0258 [-0.1356, 0.1831] | Other situations |
| lagoas.br | -0.2405 [-0.4854, -0.0665] | -0.2798 [-0.5651, -0.0048] | 0.073 [-0.0908, 0.2506] | Loss of spatially restricted species |
| lake.us | -0.2153 [-0.4433, -0.0263] | -0.1695 [-0.4234, 0.0776] | 0.093 [-0.0757, 0.2747] | Other situations |

| site_code | Δα [l-95% CI, u-95% CI] | Δγ [l-95% CI, u-95% CI] | Δβ [l-95% CI, u-95% CI] | Change in widespread or spatially restricted species |
|---|---|---|---|---|
| lancaster.uk | -0.1846 [-0.3794, 0.0152] | -0.1636 [-0.4148, 0.0872] | 0.0053 [-0.1683, 0.1695] | Other situations |
| look.us | -0.1815 [-0.38, 0.0307] | -0.1719 [-0.4231, 0.0826] | -0.0162 [-0.1977, 0.1507] | Other situations |
| marc.ar | -0.1834 [-0.3785, 0.0248] | -0.1038 [-0.3609, 0.1523] | 0.0562 [-0.1093, 0.228] | Other situations |
| mcla.us | -0.1632 [-0.3509, 0.0622] | -0.0414 [-0.3211, 0.2397] | 0.054 [-0.1063, 0.2249] | Other situations |
| moab.us | -0.1804 [-0.3769, 0.0311] | -0.0849 [-0.3485, 0.1851] | 0.0638 [-0.0966, 0.2384] | Other situations |
| msla.us | -0.1838 [-0.3788, 0.0248] | -0.1041 [-0.3631, 0.1584] | 0.0556 [-0.107, 0.2263] | Other situations |
| msla_2.us | -0.1501 [-0.3347, 0.0899] | -0.0262 [-0.3136, 0.2586] | 0.0176 [-0.1505, 0.1787] | Other situations |
| msla_3.us | -0.1707 [-0.3606, 0.0551] | -0.1199 [-0.3677, 0.136] | -0.0003 [-0.1687, 0.1624] | Other situations |
| msum.us | -0.2032 [-0.4143, -0.0121] | -0.1344 [-0.3806, 0.1163] | 0.09 [-0.0808, 0.2761] | Other situations |
| mtca.au | -0.2406 [-0.4864, -0.0683] | -0.4201 [-0.7744, -0.0744] | -0.0554 [-0.2506, 0.1167] | Loss of spatially restricted species |
| nilla.au | -0.1286 [-0.3056, 0.1346] | 0.0428 [-0.275, 0.3674] | 0.023 [-0.1441, 0.188] | Other situations |
| ping.au | -0.2343 [-0.4763, -0.0482] | -0.2589 [-0.5213, -0.0009] | 0.0724 [-0.091, 0.2491] | Loss of spatially restricted species |
| potrok.ar | -0.1573 [-0.3446, 0.0769] | -0.1116 [-0.3635, 0.1501] | -0.031 [-0.2136, 0.1391] | Other situations |
| rook.uk | -0.2072 [-0.4198, -0.0185] | -0.0478 [-0.3141, 0.2278] | 0.1817 [-0.0288, 0.4256] | Other situations |
| saana.fi | -0.2274 [-0.4557, -0.0494] | -0.337 [-0.6469, -0.0354] | -0.0192 [-0.1964, 0.1462] | Loss of spatially restricted species |
| sage.us | -0.1643 [-0.3448, 0.0652] | -0.0636 [-0.3271, 0.2027] | 0.0358 [-0.1274, 0.1971] | Other situations |
| saline.us | -0.203 [-0.4175, -0.009] | -0.1918 [-0.4408, 0.0577] | 0.0349 [-0.1288, 0.1997] | Other situations |
| sedg.us | -0.26 [-0.5265, -0.0829] | -0.6189 [-1.1423, -0.1064] | -0.1708 [-0.4567, 0.0734] | Loss of spatially restricted species |
| sereng.tz | -0.1746 [-0.3664, 0.039] | -0.1116 [-0.3689, 0.1445] | 0.0252 [-0.1423, 0.191] | Other situations |
| sevi.us | -0.1723 [-0.3658, 0.0487] | -0.172 [-0.4239, 0.0771] | -0.0342 [-0.2206, 0.1343] | Other situations |
| sgs.us | -0.1392 [-0.317, | -0.1853 [-0.4385, | -0.1543 [-0.427, | Other situations |

| site_code | Δα [l-95% CI, u-95% CI] | Δγ [l-95% CI, u-95% CI] | Δβ [l-95% CI, u-95% CI] | Change in widespread or spatially restricted species |
|---|---|---|---|---|
| | 0.1092] | 0.0647] | 0.0818] | |
| shps.us | -0.2123 [-0.4319, -0.0174] | -0.253 [-0.5209, 0.0051] | 0.0137 [-0.1593, 0.1798] | Other situations |
| sier.us | -0.1125 [-0.2864, 0.1655] | 0.1126 [-0.2536, 0.4853] | 0.0333 [-0.1335, 0.198] | Other situations |
| smith.us | -0.1754 [-0.3656, 0.0435] | -0.1661 [-0.4131, 0.0835] | -0.0221 [-0.1992, 0.1432] | Other situations |
| spin.us | -0.217 [-0.4405, -0.0307] | -0.1363 [-0.39, 0.1178] | 0.131 [-0.0485, 0.3353] | Other situations |
| sval.no | -0.1567 [-0.3434, 0.0771] | -0.0415 [-0.3041, 0.2327] | 0.033 [-0.1319, 0.1987] | Other situations |
| temple.us | -0.2157 [-0.4384, -0.0235] | -0.2568 [-0.5186, 0.0019] | 0.0159 [-0.1504, 0.1824] | Other situations |
| thth.is | -0.194 [-0.402, 0.0034] | -0.1157 [-0.3701, 0.1447] | 0.0824 [-0.0821, 0.259] | Other situations |
| tmlr.is | -0.1677 [-0.36, 0.0535] | -0.1196 [-0.3681, 0.136] | -0.0005 [-0.1752, 0.1639] | Other situations |
| trel.us | -0.1992 [-0.4047, -0.0041] | -0.1906 [-0.4477, 0.0595] | 0.0225 [-0.1443, 0.189] | Other situations |
| ukul.za | -0.1777 [-0.3676, 0.038] | -0.1006 [-0.3562, 0.1626] | 0.0406 [-0.122, 0.2095] | Other situations |
| unc.us | -0.253 [-0.5182, -0.0774] | -0.4709 [-0.8686, -0.082] | -0.0596 [-0.2601, 0.1169] | Loss of spatially restricted species |
| valm.ch | -0.2081 [-0.423, -0.014] | -0.2233 [-0.4889, 0.0354] | 0.0258 [-0.1414, 0.1936] | Other situations |
| veluwe.nl | -0.1485 [-0.3297, 0.0946] | -0.0172 [-0.2972, 0.2769] | 0.0233 [-0.1452, 0.1843] | Other situations |
| yarra.au | -0.2078 [-0.4271, -0.0132] | -0.3555 [-0.6704, -0.0414] | -0.0961 [-0.3189, 0.0981] | Loss of spatially restricted species |

**Table S5.**

**Summary of model fitting for the effects of nutrient addition on alpha, gamma, and beta diversity ($\Delta\alpha$, $\Delta\gamma$, and $\Delta\beta$) for native and non-native species four years after treatment began.** We calculated $\alpha$ and $\gamma$ diversity using species richness. We fitted the random intercept models using the brm function. For $\Delta\alpha$, $\Delta\gamma$, and $\Delta\beta$, we included random intercepts among sites, model was coded as: diversity (log scale) ~ 1 + (1 |sites). Bulk-ESS is the effective sample size, is useful as a diagnostic for the sampling efficiency. Tail_ESS is the minimum of the effective sample sizes for 5% and 95% quantiles. Low Rhats indicates model convergence were achieved. All models used 1000 iterations for warmup and 3000 iterations

| Species groups | Diversity facets | Estimate | l-95% CI | u-95% CI | Rhat | Bulk_ESS | Tail_ESS | Number of sites |
|---|---|---|---|---|---|---|---|---|
| Native | $\Delta\alpha$ | -0.2118 | -0.2861 | -0.1374 | 1.0009 | 12510.4909 | 8656.293 | 69 |
| Native | $\Delta\beta$ | 0.0005 | -0.0548 | 0.0552 | 1.0172 | 342.3685 | 806.045 | 69 |
| Native | $\Delta\gamma$ | -0.2113 | -0.3007 | -0.1234 | 1.0033 | 2001.1538 | 3930.769 | 69 |
| Non-native | $\Delta\alpha$ | -0.1155 | -0.2278 | -0.0038 | 1.0007 | 9704.4354 | 8918.270 | 42 |
| Non-native | $\Delta\beta$ | 0.0454 | -0.0466 | 0.1380 | 1.0020 | 2009.6608 | 1466.581 | 42 |
| Non-native | $\Delta\gamma$ | -0.0499 | -0.1972 | 0.1015 | 1.0046 | 1921.3336 | 2824.314 | 42 |

**Table S6.**

**Summary of model fitting for the effects of nutrient addition on alpha, gamma, and beta diversity (Δα, Δγ, and Δβ) for different life forms four years after treatment began.** We calculated α and γ diversity using species richness. We fitted the random intercept models using the brm function. For Δα, Δγ, and Δβ, we included random intercepts among sites, model was coded as: diversity (log scale) ~ 1 + (1 |sites). Bulk-ESS is the effective sample size, is useful as a diagnostic for the sampling efficiency. Tail_ESS is the minimum of the effective sample sizes for 5% and 95% quantiles. Low Rhats indicates model convergence were achieved. All models used 1000 iterations for warmup and 3000 iterations

| Species groups | Diversity facets | Estimate | l-95% CI | u-95% CI | Rhat | Bulk_ESS | Tail_ESS | Number of sites |
|---|---|---|---|---|---|---|---|---|
| FORB | Δα | -0.1941 | -0.2908 | -0.0939 | 1.0009 | 10299.5121 | 8854.6421 | 68 |
| FORB | Δβ | 0.0854 | -0.0155 | 0.1874 | 1.0054 | 1579.1276 | 3469.0268 | 68 |
| FORB | Δγ | -0.1139 | -0.2223 | -0.0125 | 1.0254 | 191.9344 | 4084.4711 | 68 |
| GRAMINOID | Δα | -0.1320 | -0.1998 | -0.0628 | 1.0005 | 10726.0783 | 8985.3000 | 72 |
| GRAMINOID | Δβ | 0.0061 | -0.0393 | 0.0546 | 1.0038 | 1644.1281 | 1257.2676 | 72 |
| GRAMINOID | Δγ | -0.1276 | -0.2091 | -0.0476 | 1.0060 | 2455.7868 | 4956.4376 | 72 |
| LEGUME | Δα | -0.1894 | -0.3169 | -0.0600 | 1.0010 | 12678.7708 | 8585.2324 | 34 |
| LEGUME | Δβ | 0.0049 | -0.1681 | 0.1759 | 1.0009 | 3779.7706 | 4843.1788 | 34 |
| LEGUME | Δγ | -0.1887 | -0.3895 | -0.0029 | 1.0095 | 703.4521 | 708.2061 | 34 |
| WOODY | Δα | -0.1828 | -0.3425 | -0.0197 | 1.0005 | 8915.9640 | 7764.9732 | 23 |
| WOODY | Δβ | -0.1426 | -0.2950 | 0.0027 | 1.0016 | 2747.1866 | 3021.2355 | 23 |
| WOODY | Δγ | -0.2887 | -0.5220 | -0.0563 | 1.0027 | 1919.7104 | 2497.4429 | 23 |

**Table S7.**

**Principal investigators contributing data but are not authors; Names match those in Table S1. Their effort in providing data is critical to this manuscript.**

| site_code | PI name | Institution |
|---|---|---|
| ahth.is | Isabel Barrio | University of Iceland |
| amlr.is | Isabel Barrio | University of Iceland |
| arch.us | Elizabeth Boughton | MacArthur Agro-ecology Research Center |
| azi.cn | Chengjin Chu | Lanzhou University |
| azi.cn | Guozhen Du | Lanzhou University |
| azi.cn | Qi Li | Northwest Institute of Plateau Biology |
| azi.cn | Wei Li | Iowa State University |
| azi.cn | Gang Wen | Lanzhou University |
| azitwo.cn | Guozhen Du | Lanzhou University |
| azitwo.cn | Zhengwei Ren | Lanzhou University |
| badlau.de | Julia Siebert | German Centre for Integrative Biodiversity Research (iDiv) |
| barta.us | David Wedin | University of Nebraska, Lincoln |
| bayr.de | Marie Spohn | NULL |
| bnbt.us | Brent Mortensen | Benedictine College |
| bnbt.us | Janet Paper | Benedictine College |
| bogong.au | Joslin Moore | Victoria State Government |
| burrawan.au | Jennifer Firn | Queensland University of Technology |
| burren.ie | Yvonne Buckley | Trinity College Dublin, the University of Dublin |
| cbgb.us | Lori Biederman | Iowa State University |
| cbgb.us | Kirsten Hofmockel | Iowa State University |
| cbgb.us | Lauren Sullivan | Iowa State University |
| cdcr.us | Adam Kay | University of St. Thomas |
| chilcas.ar | Enrique Chaneton | Universidad de Buenos Aires |
| chilcas.ar | Laura Yahdjian | Universidad de Buenos Aires |
| comp.pt | Maria Caldeira | Technical University of Lisbon |
| elliot.us | Elsa Cleland | University of California, San Diego |
| frue.ch | Sabine Güsewell | ETH Zurich |
| frue.ch | Andy Hector | University of Zurich |
| hall.us | Rebecca McCulley | University of Kentucky |
| hall.us | Jim Nelson | University of Kentucky |
| hart.us | Nicole DeCrappeo | USGS |
| hart.us | David Pyke | USGS |
| hero.uk | Mick Crawley | Imperial College at Silwood Park |
| jena.de | Anne Ebeling | Friedrich-Schiller-Universität Jena |
| kbs.us | Lars Brudvig | Michigan State University |
| kibber.in | Mahesh Sankaran | National Centre for Biological Sciences |
| kidman.au | Anna Richards | CSIRO |
| koffler.ca | Arthur Weiss | University of Toronto Scarborough |

| site_code | PI name | Institution |
|---|---|---|
| konz.us | Kimberly Komatsu | University of North Carolina - Greensboro |
| konz.us | Melinda Smith | Colorado State University |
| lagoas.br | Lucíola Lannes | Universidade Estadual Paulista - UNESP |
| lake.us | Lori Biederman | Iowa State University |
| marc.ar | Juan Alberti | Universidad Nacional de Mar del Plata |
| moab.us | Brooke Osborne | US Geological Survey |
| moab.us | Sasha Reed | US Geological Survey |
| msla.us | Mary Ellyn DuPre | MPG Ranch |
| msla.us | Kelly Laflamme | MPG Ranch |
| msla.us | Yiva Lekberg | MPG Ranch |
| msla_2.us | Mary Ellyn DuPre | MPG Ranch |
| msla_2.us | Yiva Lekberg | MPG Ranch |
| msla_3.us | Mary Ellyn DuPre | MPG Ranch |
| msla_3.us | Yiva Lekberg | MPG Ranch |
| msum.us | Alison Wallace | Minnesota State University Moorhead |
| mtca.au | Suzanne Prober | CSIRO |
| ping.au | Jodi Price | The University of Western Australia |
| ping.au | Rachel Standish | The University of Western Australia |
| potrok.ar | Hector Bahamonde | UNPA - CONICET |
| potrok.ar | Pablo Peri | UNPA - CONICET |
| rook.uk | Mick Crawley | Imperial College at Silwood Park |
| sage.us | Louie Yang | University of California, Davis |
| saline.us | Kimberly Komatsu | University of North Carolina - Greensboro |
| saline.us | Melinda Smith | Colorado State University |
| sereng.tz | T. Anderson | Wake Forest University |
| sevi.us | Scott Collins | University of New Mexico |
| sevi.us | Laura Ladwig | University of New Mexico |
| sgs.us | Dana Blumenthal | USDA-ARS |
| sgs.us | Cynthia Brown | Colorado State University |
| sgs.us | Julia Klein | Colorado State University |
| sgs.us | Alan Knapp | Colorado State University |
| smith.us | Janneke Hille Ris Lambers | University of Washington |
| spin.us | Rebecca McCulley | University of Kentucky |
| spin.us | Jim Nelson | University of Kentucky |
| temple.us | Philip Fay | USDA ARS |
| temple.us | Jason Martina | Texas State University |
| thth.is | Isabel Barrio | University of Iceland |
| tmlr.is | Isabel Barrio | University of Iceland |
| trel.us | Andrew Leakey | University of Illinois at Urbana-Champaign |
| ukul.za | Kevin Kirkman | University of KwaZulu-Natal |
| unc.us | Charles Mitchell | University of North Carolina |
| unc.us | Justin Wright | Duke University |
| yarra.au | Raul Ochoa Hueso | University of Western Sydney |

**Table S8.**
**Contributions of each author to the manuscript.** Author contributions to individual tasks are marked with x.

| Full name | Sites used in analysis | Developed and framed research questions | Analyzed data | Contributed to data analyses | Wrote the paper | Contributed to paper writing | Site coordinator | Nutrient Network coordinator | Site level acknowledgments |
|---|---|---|---|---|---|---|---|---|---|
| Qingqing Chen | | x | x | | x | | | | |
| Shane Blowes | | x | | x | | x | | | |
| Emma Ladouceur | | x | | x | | x | | | |
| W. Stanley Harpole | hopl.us, mcla.us, sier.us, cbgb.us | | | | | x | x | x | |
| Jonathan Chase | | x | | | | x | | | |
| Jonathan D. Bakker | smith.us | | | x | | x | x | | |
| Nico Eisenhauer | badlau.de | | | | | x | x | | NE acknowledges funding by the German Research Foundation (DFG, FZT 118, 202548816; Ei 862/29-1). |
| Eric W. Seabloom | bnch.us, look.us, hopl.us, mcla.us, sier.us, cdcr.us | | | | | x | x | x | |
| Pablo L. Peri | potrok.ar | | | | | x | x | | |
| Pedro M. Tognetti | chilcas.ar | | | | | x | x | | |
| Risto Virtanen | kilp.fi,saana.fi | | | | | x | x | | |
| Sally A. Power | Yarra.au | | | | | x | x | | |

| Full name | Sites used in analysis | Developed and framed research questions | Analyzed data | Contributed to data analyses | Wrote the paper | Contributed to paper writing | Site coordinator | Nutrient Network coordinator | Site level acknowledgments |
|---|---|---|---|---|---|---|---|---|---|
| George R. Wheeler | cdpt.us | | | | | x | x | | |
| Michelle Tedder | ukul.za, gilbza | | | | | x | x | | |
| Anita C. Risch | valm.ch | | | | | x | x | | |
| Sumanta Bagchi | kibber.in | | | | | | | | |
| Jane A. Catford | nilla.au | | | | | x | x | | |
| Carly J. Stevens | lancaster.uk | | | | | | | | |
| Christiane Roscher | jena.de | | | | | x | x | | |
| Yujie Niu | bayr.de | | | | | x | | | |
| Sylvia Haider | badlau.de | | | | | x | x | | |
| Maria C. Caldeira | comp.pt | | | | | x | x | | to Companhia das Lezirias for granting access to the study site and Portuguese Science Foundation (FCT) for Forest Research Centre funding (UIDB/00239/2020). |
| Peter B. Adler | shps.us | | | | | x | x | | |
| Jason P. Martina | temple.us | | | | | x | x | | |
| Miguel N. Bugalho | comp.pt | | | | | x | x | | |
| Johannes M H Knops | cdpt.us | | | | | x | x | | |
| Elizabeth T. Borer | bnch.us, look.us, hopl.us, mcla.us, sier.us, cdcr.us | | | | | x | x | x | |

| Full name | Sites used in analysis | Developed and framed research questions | Analyzed data | Contributed to data analyses | Wrote the paper | Contributed to paper writing | Site coordinator | Nutrient Network coordinator | Site level acknowledgments |
|---|---|---|---|---|---|---|---|---|---|
| Christopher R. Dickman | ethamc.au, ethass.au | | | | | x | x | | |
| Anke Jentsch | bayr.de | | | | | x | x | | AJ acknowledges funding by the Federal Ministry of Education and Research (BMBF) grant 031B0516C and by the Upper Franconian Trust Oberfrankenstiftung grant FP00237 |
| Yann Hautier | frue.ch | | | | | x | x | | |
| Andrew MacDougall | Cowi.ca | | | | | x | x | | |
| John W. Morgan | kiny.au, bogong.au | | | | | x | x | | |
| Glenda M. Wardle | ethamc.au, ethass.au | | | | | x | x | | GMW acknowledges Bush Heritage Australia and the Wangkamadla people for access to the sites and the ARC, and TERN funded by NCRIS for funding |
| Pedro Daleo | marc.ar | | | | | x | x | | |
| Catalina Estrada | hero.uk, rook.uk | | | | | x | x | | Imperial College London, Department of Life Sciences |
| Ian Donohue | burren.ie | | | | | x | x | | |
| G.F. (Ciska) Veen | veluwe.nl | | | | | x | x | | |
| Daniel S Gruner | sage.us | | | | | x | x | | |
| Harry Olde Venterink | lagoas.br | | | | | x | x | | |
| Erika Hersch-Green | | | | | | x | | | |
| Anu Eskelinen | kilp.fi | | | | | x | x | | |

| Full name | Sites used in analysis | Developed and framed research questions | Analyzed data | Contributed to data analyses | Wrote the paper | Contributed to paper writing | Site coordinator | Nutrient Network coordinator | Site level acknowledgments |
|---|---|---|---|---|---|---|---|---|---|
| Nicole Hagenah | ukul.za | | | | | x | x | | |
| Carla D'Antonio | sedg.us | | | | | x | x | | |
| Marc W. Cadotte | Koffler | | | | | x | x | | |
| Brooke B. Osborne | moab.us | | | | | x | x | | |
| Lauri Laanisto | sval.no | | | | | x | x | | Funded by Estonian Academy of Sciences (research professorship for Arctic studies) |
| Petr Macek | sval.no | | | | | x | x | | Funded by Estonian Academy of Sciences (research professorship for Arctic studies) |
